# Glioblastoma stem cells induce quiescence in surrounding neural stem cells via Notch signalling

**DOI:** 10.1101/856062

**Authors:** Katerina Lawlor, Maria Angeles Marques-Torrejon, Gopuraja Dharmalingham, Yasmine El-Azhar, Michael D. Schneider, Steven M. Pollard, Tristan A. Rodríguez

## Abstract

There is increasing evidence suggesting that adult neural stem cells (NSCs) are a cell of origin of glioblastoma, the most aggressive form of malignant glioma. The earliest stages of hyperplasia are not easy to explore, but likely involve a cross-talk between normal and transformed NSCs. How normal cells respond to this cross-talk and if they expand or are outcompeted is poorly understood. Here we have analysed the interaction of transformed and wild-type NSCs isolated from the adult mouse subventricular zone neural stem cell niche. We find that transformed NSCs are refractory to quiescence-inducing signals. Unexpectedly, however, we also demonstrate that these cells induce a quiescent-like state in surrounding wild-type NSC. We find that this response is cell-cell contact-dependent and that transformed cells activate the Notch pathway in adjacent wild-type NSCs, an event that stimulates their entry into quiescence. Our findings therefore suggest that oncogenic mutations may be propagated in the stem cell niche not just though cell-intrinsic advantages, but also by outcompeting neighbouring stem cells through signalling repression of their proliferation.

## Introduction

In the adult mammalian brain, neural stem cells (NSCs) can be primarily found in specific neurogenic regions. In rodents these are the subgranular zone (SGZ) of the dentate gyrus (DG) and the subventricular zone (SVZ), which lines the lateral ventricles (Doetsch et al., 1999; Quinones-Hinojosa et al., 2006; Sanai et al., 2004). In the SVZ there are both quiescent NSCs and activated NSCs that coexist in the niche (Quinones-Hinojosa et al., 2006). The quiescent NSC subpopulation make up the majority of NSCs in the adult SVZ and are capable of completely regenerating this niche following anti-mitotic drug treatment (Doetsch et al., 1999; Giachino and Taylor, 2009; Pastrana et al., 2009). Tight regulation of the switch between these two states is vital to ensure that the pool of NSCs does not accumulate DNA damage or become exhausted over time and therefore understanding how this is achieved is of great interest. One key event in NSC activation is the upregulation of EGFR, which is highly expressed in activated NSCs and progenitor cells but absent in quiescent NSCs. Thus, cell surface expression of EGFR is often used as a marker to differentiate these two populations (Codega et al., 2014; Pastrana et al., 2009). What feedback mechanisms exist between activated EGFR^+ve^ NSCs and quiescent NSCs to ensure stem cell homeostasis is poorly understood.

Over the last couple of decades, evidence has accumulated to suggest that adult NSCs are a cell of origin in certain brain tumours. One possibility is that these cells accumulate driver mutations over time, which compromise the normal controls on their proliferation and migration. These transformed NSCs could then escape the niche and acquire further mutations resulting in tumour formation. This has been hypothesised to be the case in glioblastoma (GBM), the most aggressive subtype of malignant glioma (Louis et al., 2007; Rispoli et al., 2014). In particular, the discovery of a subpopulation of cells within GBM tumours with stem cell characteristics, known as glioblastoma stem cells (GSCs), has lent weight to this notion. This GSC population in GBM tumours share many common features with adult NSCs, including expression of stem and progenitor markers, such as CD133, SOX2 and Nestin, self-renewal capacity, and the ability to generate multilineage progeny (Cheng et al., 2013; Denysenko et al., 2010; Singh et al., 2003; Singh et al., 2004; Stieber et al., 2014; Zhang et al., 2006). Most importantly, GSCs are also able to generate tumours in mice which recapitulate all the classical features of GBM, even when injected in very small numbers (Galli et al., 2004; Singh et al., 2004). For this reason, they are sometimes also referred to as brain tumour initiating cells.

Adult NSCs can be isolated from the SVZ of murine brains and cultured as 3D neurospheres or adherent cultures under conditions that promote symmetric self-renewal and prevent differentiation (Conti et al., 2005; Pollard et al., 2006; Pollard et al., 2009). Furthermore, by using NSCs engineered with oncogenic drivers, these *in vitro* culture systems can also be useful to model brain tumour development and cross-comparison with wild-type counterparts. Here we exploit this system to understand how transforming mutations affect the quiescent or activation status of NSCs. We find that transformed NSCs are refractory to quiescent inducing signals. Importantly, we also demonstrate that transformed NSCs signal via Notch and RBPJ to induce a quiescent-like state in surrounding wild-type NSCs. These results therefore provide new insight into how oncogenic mutations affect not only the proliferative potential of transformed cells, but also provide them with a competitive advantage in the niche by suppressing the proliferation of surrounding cells.

## Results

### Transformed NSCs are refractory to quiescence-inducing signals

To understand how transforming mutations affect the quiescent or activation status of NSCs we compared NSCs derived from wild-type mice to those isolated from *Ink4a/Arf*^*-/-*^ mice and transduced with a retrovirus expressing the EGFRvIII mutation (Bruggeman et al., 2007). This combination of mutations is frequently observed in human GBM (Crespo et al., 2015) and induces transformation in NSCs, as indicated by their ability to generate tumours *in vivo* that recapitulate many of the features of human GBM tumours (Bruggeman et al., 2007; Marques-Torrejon et al., 2018). We first compared the proliferative potential of both these cell types. We observed that, when grown in conditions that promote the activated/proliferative NSC state (Pollard et al., 2006), transformed NCSs displayed a small growth advantage (Figure 1A). Analysis of the prevalence of BrdU incorporation revealed that this difference was most likely accounted for by a small difference in the rate of cell division at day four of culture, when transformed cells divided faster (Supplementary Figure 1A-B). However, overall these results indicate that the pattern of proliferation of wild-type and *Ink4a/Arf*^*-/-*^; EGFRvIII NSCs (IE-NSCs) in growth-promoting conditions was remarkably similar.

**Figure 1.**
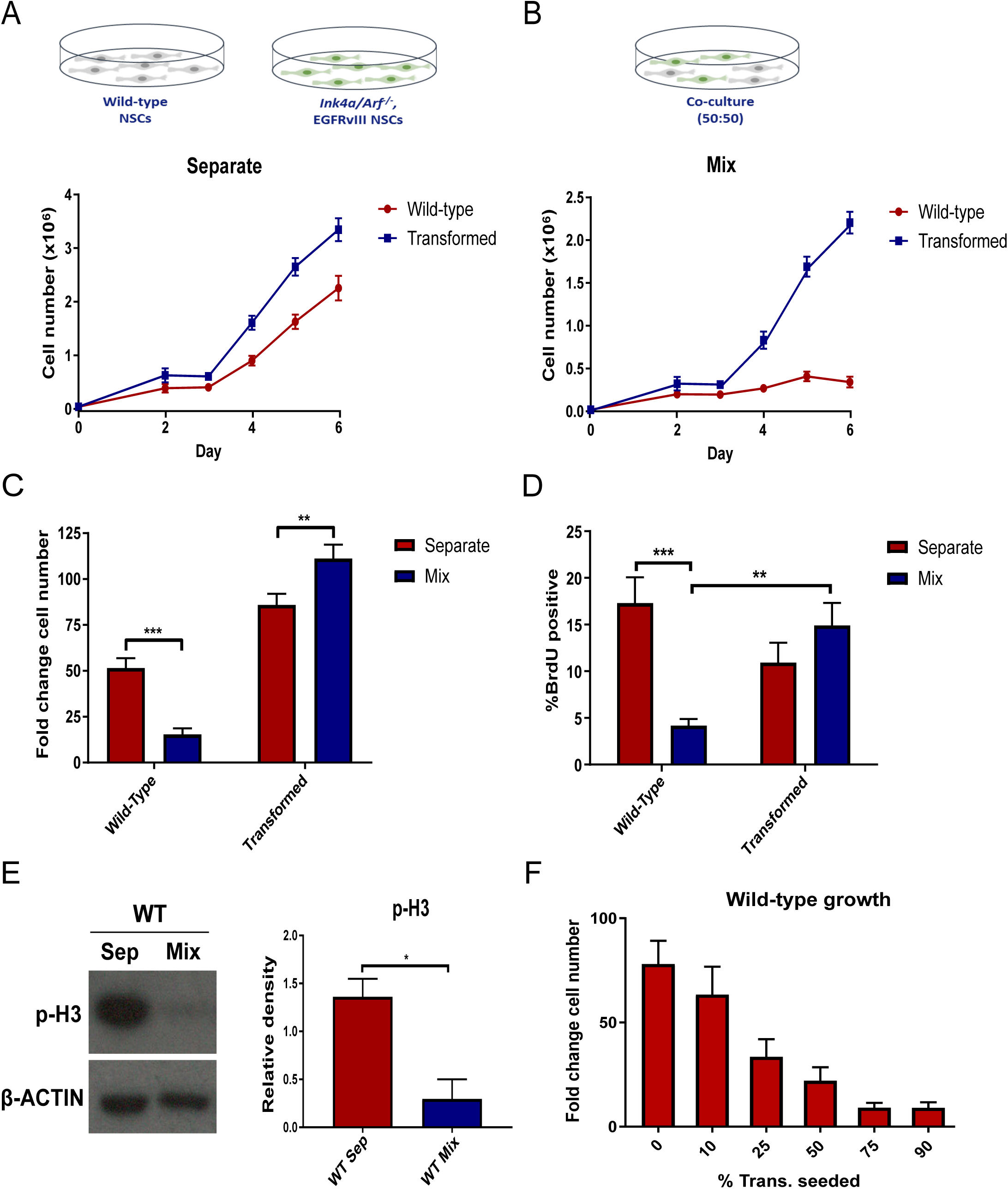
WT NSCs show reduced proliferation in the presence of transformed NSCs. **A** Growth curves of WT and transformed IE NSCs cultured separately over six days. **B** Growth curves of wild-type and transformed NSCs in a 50:50 co-culture. **C** Quantification of fold change in cell number relative to seeding density for each culture condition. WT NSCs dramatically reduce their proliferation while transformed cells increase their proliferation relative to separate culture. N>10, ANOVA followed by Sidak’s multiple comparisons test. **D** Proportion of BrdU-positive cells in each culture condition at day five following a two-hour BrdU chase. WT NSCs show greatly reduced BrdU incorporation in co-culture. N=10, ANOVA followed by Sidak’s multiple comparisons test. **E** Representative western blot for phospho-histone-3 (p-H3) in WT NSCs sorted at day five and quantification of p-H3 density relative to β-actin. p-H3 expression is markedly reduced in co-cultured WT NSCs. N=3, Student’s paired T test. **F** Fold change cell number of WT NSCs cultured in co-cultures with varying proportions of transformed NSCs. WT NSCs show smaller increases in cell number in co-cultures with a greater proportion of transformed NSCs. N=3.

We next investigated if IE-NSCs have a similar response to quiescence-inducing signals when compared to wild-type NSCs. BMP signalling is known to induce a reversible quiescent-like state in hippocampal and embryonic stem cell-derived NSCs (Martynoga et al., 2013; Mira et al., 2010) and the BMP antagonist Noggin enhances NSC expansion both *in vitro* and *in vivo* (Lim et al., 2000; Mira et al., 2010; Yousef et al., 2015). For this reason, we treated wild-type and transformed NSCs with 50ng/ml BMP4. We observed that, although this induced similar levels of SMAD1/5/8 phosphorylation in both cell types (Supplementary Figure 1C) and efficiently suppressed the proliferation of wild-type cells, it had little effect on the growth of the transformed cells (Supplementary Figure 1D). This suggests that IE-NSCs cells are refractory to the induction of quiescence by BMP signalling activation.

### Transformed NSCs suppress the proliferation of surrounding NSCs

The experiments described above allowed us to test the proliferative potential of wild-type and transformed cells as isolated cell populations. However, we postulated that in the niche these cell types are likely to co-exist. For this reason, we next studied the proliferation patterns of both cell types in a 50:50 co-culture and compared these proliferative patterns to the growth of the cells in homotypic (separate) cultures. IE-NSCs were GFP-labelled, allowing us to quantify the relative cell proliferation rates in co-culture using flow cytometry. Unexpectedly, we observed that, in contrast to the exponential growth observed for both cell types in separate culture, in co-culture wild-type NSCs showed a significant growth arrest (Figure 1B). Transformed NSCs therefore do not facilitate normal NSC growth as might be anticipated; rather, they deploy some active mechanism to suppress their expansion and outcompete them.

Quantification of the fold change in cell number indicated that the wild-type NSCs increased their cell number only around 15-fold over six days in co-culture, compared to more than 50-fold when cultured separately (Figure 1C). This dramatic reduction in the proliferation of wild-type NSCs in the co-culture condition was also confirmed using BrdU incorporation assays and quantifying expression mitotic marker phospho-Histone3 (Figure 1D-E and Supplementary Figure 1C-D). These results suggest that IE-NSCs have an inhibitory effect on the proliferation of wild-type NSCs. To test this possibility further, co-cultures were established with varying proportions of each cell type, ranging from 10 to 90%, and the total number of cells present in co-culture was kept constant. Analysis of the fold change in wild-type NSC number revealed that the reduction in their proliferation was directly proportional to the percentage of transformed NSCs present (Figure 1F), to the extent that, when 75% of the cells in co-culture were transformed, the wild-type NSCs barely increased in number during the 6-day experiment. Together, these results indicate that IE-NSCs suppress the proliferation of wild-type NSCs.

### Transformed cells induce a quiescent-like state in wild-type NSCs

To gain a deeper insight into the mechanism of proliferative arrest of co-cultured wild-type NSCs, we compared their transcriptional profile to wild-type NSCs grown in separate culture. Cells were isolated by FACS after 5 days of co-culture, and RNA-seq was performed. Analysis of the differential gene expression yielded 1441 differentially expressed genes that were then studied by Ingenuity Pathway Analysis, which uses functional annotations and interactions of genes to identify enriched pathways and upstream regulators. It was immediately apparent that the majority of the top canonical pathways that were enriched in this comparison were related to cell cycle control and DNA repair (Figure 2A). For example, the top enriched pathway was ‘cell cycle control of chromosomal replication’, which includes important cell cycle-associated genes, such as multiple CDKs, MCM proteins and PCNA. The genes in this pathway were broadly downregulated in wild-type NSCs in co-culture compared to separately cultured cells (Figure 2B), consistent with a proliferation arrest when surrounded by transformed cells.

**Figure 2.**
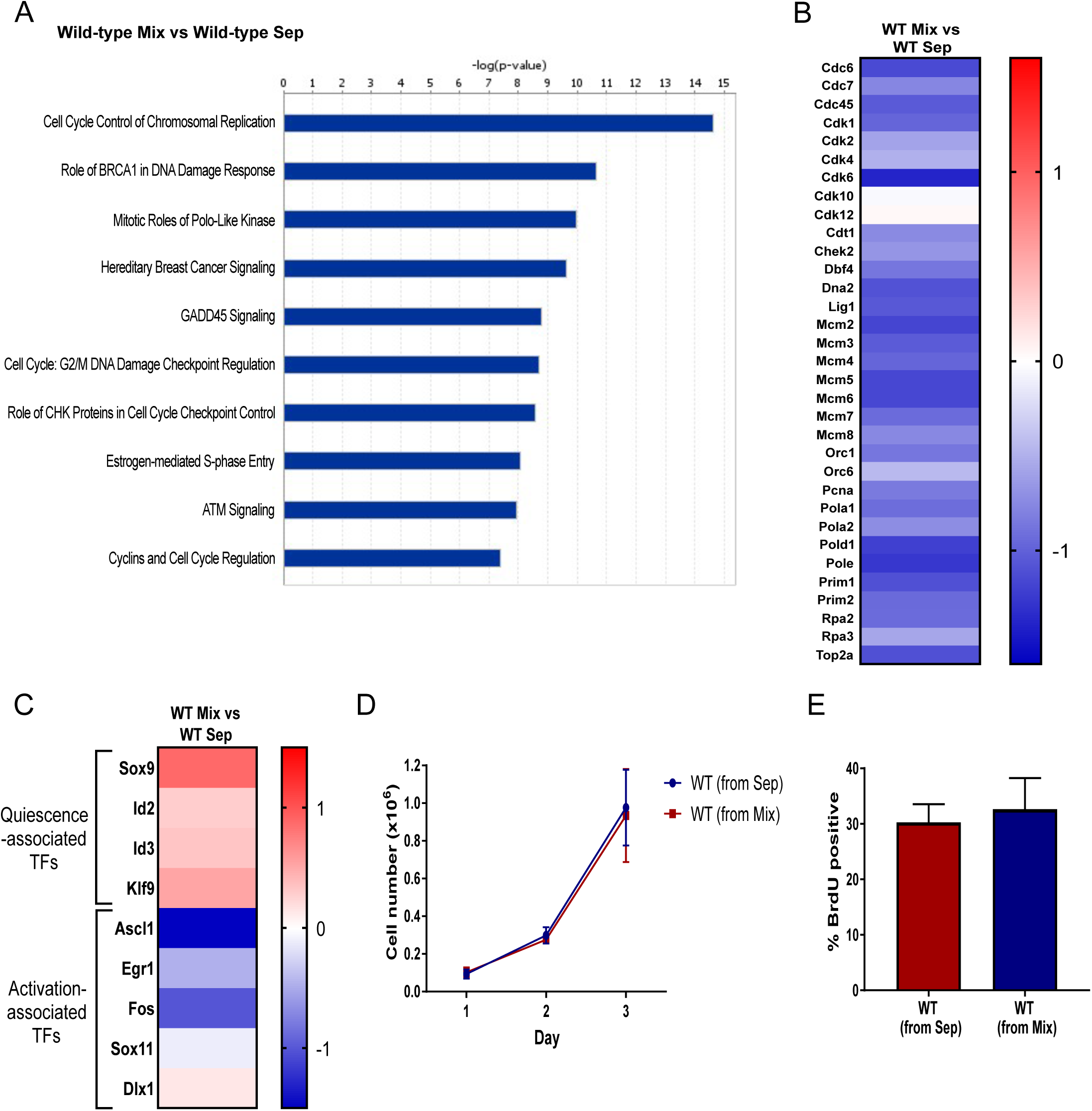
Wild-type NSCs adopt a quiescent phenotype in the presence of transformed NSCs. **A** RNA sequencing was performed on NSCs sorted by FACS following five days’ culture in either homogenous or co-culture conditions. Differential gene expression analysis was performed on these samples from three independent experiments. The top canonical pathways enriched in the comparison of WT NSCs in separate or mixed culture. Pathways related to cell cycle and DNA repair are highly enriched. **B** Heat map showing the fold change values for all genes in the ‘Cell Cycle Control of Chromosomal Replication’ pathway for this comparison with an adjusted p-value <0.1. **C** Heat map showing the change in gene expression of certain transcription factors (TFs) between the WT NSCs cultured separately or in co-culture. These transcription factors have been particularly highlighted by transcriptional profiling studies that sought to compare quiescent and activated NSCs. Quiescence-associated genes were broadly upregulated in the co-cultured NSCs, while activation-associated genes were broadly downregulated. **D** Growth curves showing cell numbers for three days following replating. **E** Proportion of BrdU-positive cells at day three after replating. No difference could be found in the growth of WT NSCs which had previously been co-cultured compared to cells grown separately. N=3. **D** and **E** WT NSCs were cultured for 5 days either in co-culture or separately and then sorted by FACS and replated at 100,000 cells/well.

One possible explanation for this cell cycle arrest is differentiation into a post-mitotic cell types, such as immature neurons. The expression of NSC markers, together with markers for intermediate progenitors and differentiated cell types (Zhang and Jiao, 2015), was therefore explored in our RNA-seq data-set. No broad downregulation of classical NSC markers, such as *Nestin, Gfap* and *Pax6* was observed in co-cultured wild-type NSCs compared to their separate counterparts. Likewise, markers of intermediate or differentiated cell types, such as *Doublecortin, Olig2, S100b* or *Tubb3*, were not upregulated (Supplementary Figure 2A), indicating there was no increase in differentiation, an observation that was consistent with the morphology of these cells that was not that of oligodentrocyte progenitor cells, oligodentrocytes or neurons. Interestingly, we did observe an upregulation in the expression of the glial markers, *Gfap, Glt1* and *Glast*, which are enriched in quiescent radial glia-like NSCs (Codega et al., 2014; Llorens-Bobadilla et al., 2015). These observations suggest that wild-type NSCs are not differentiating, but instead may be driven into a quiescent state.

To explore this last possibility we analysed our RNA-seq data-set for the expression of a broader set of quiescent and activated NSC markers. We found that the quiescence-associated transcriptional regulators, *Sox9, Id2, Id3* and *Klf9*, which are upregulated in quiescent SVZ NSCs (Llorens-Bobadilla et al., 2015; Morizur et al., 2018), were all also upregulated in the co-cultured wild-type NSCs (Figure 2C). We also observed that the markers of activated/proliferating NSCs *Ascl1, Egr1, Fos* and *Sox11* (Andersen et al., 2014; Llorens-Bobadilla et al., 2015; Morizur et al., 2018) were downregulated in these cells. When we compared our RNA-seq data-set with a list of transcription factors and co-factors that have been found to be differentially expressed between activated and quiescent SVZ NSCs (Morizur et al., 2018), we found that of the 14 genes enriched in quiescent cells in this study, 10 were upregulated in co-cultured wild-type NSCs, and of the 61 genes enriched in activated NSCs, 50 were downregulated (Supplementary Figure 2B). Together these findings therefore support the hypothesis that the co-cultured NSCs are adopting a quiescent phenotype rather than becoming senescent.

A key feature of a quiescent phenotype is its reversibility. To test if the proliferation arrest of wild-type NSCs in co-culture is reversible, we sorted wild-type and transformed cells after 5 days in co-culture and re-plated them. In parallel, wild-type NSCs that had been cultured separately were mixed with transformed cells just before sorting and then re-plated. Importantly, we observed that previously co-cultured wild-type NSCs displayed a similar proliferation rate to NSCs that had been separately cultured throughout, as judged by their growth curves and rate of BrdU incorporation (Figure 2E-F). The reversible nature of the proliferation arrest of co-cultured wild-type NSCs provides further evidence that they are entering a quiescent state.

### Cell contact is required for the inhibitory effect of transformed cells

To understand the mechanisms underlying the interaction between wild-type and transformed NSCs, we first analysed the possible involvement of the mTOR signalling pathway. mTOR is a metabolic regulator that senses growth factor and nutrient inputs, is activated in transit amplifying progenitor cells and its inhibition induces a reversible quiescence-like phenotype in adult NSCs (Paliouras et al., 2012). Analysis of the expression of ribosomal protein S6 phosphorylation (p-S6), a read-out of mTOR activity indicated that mTOR signalling levels were reduced in the co-cultured wild-type NSCs, not only relative to these same cells in separate culture, but also to transformed NSCs in co-culture (Figure 3A). We next tested if constitutive mTOR activation was sufficient to prevent these wild-type NSCs from entering quiescence in co-culture. For this we mutated *Tsc2*, an mTOR repressor in wild-type NSCs, using CRISPR-Cas9 targeting (Supplementary Figure 2C). However, we found that, although this was sufficient to sustain strong mTOR pathway activation, *Tsc2*^-/-^ NSCs still entered a proliferation arrest when co-cultured with transformed NSCs (Supplementary Figure 2D-F). This indicates that mTOR inhibition is not the primary reason for the quiescent phenotype of co-cultured wild-type NSCs.

**Figure 3.**
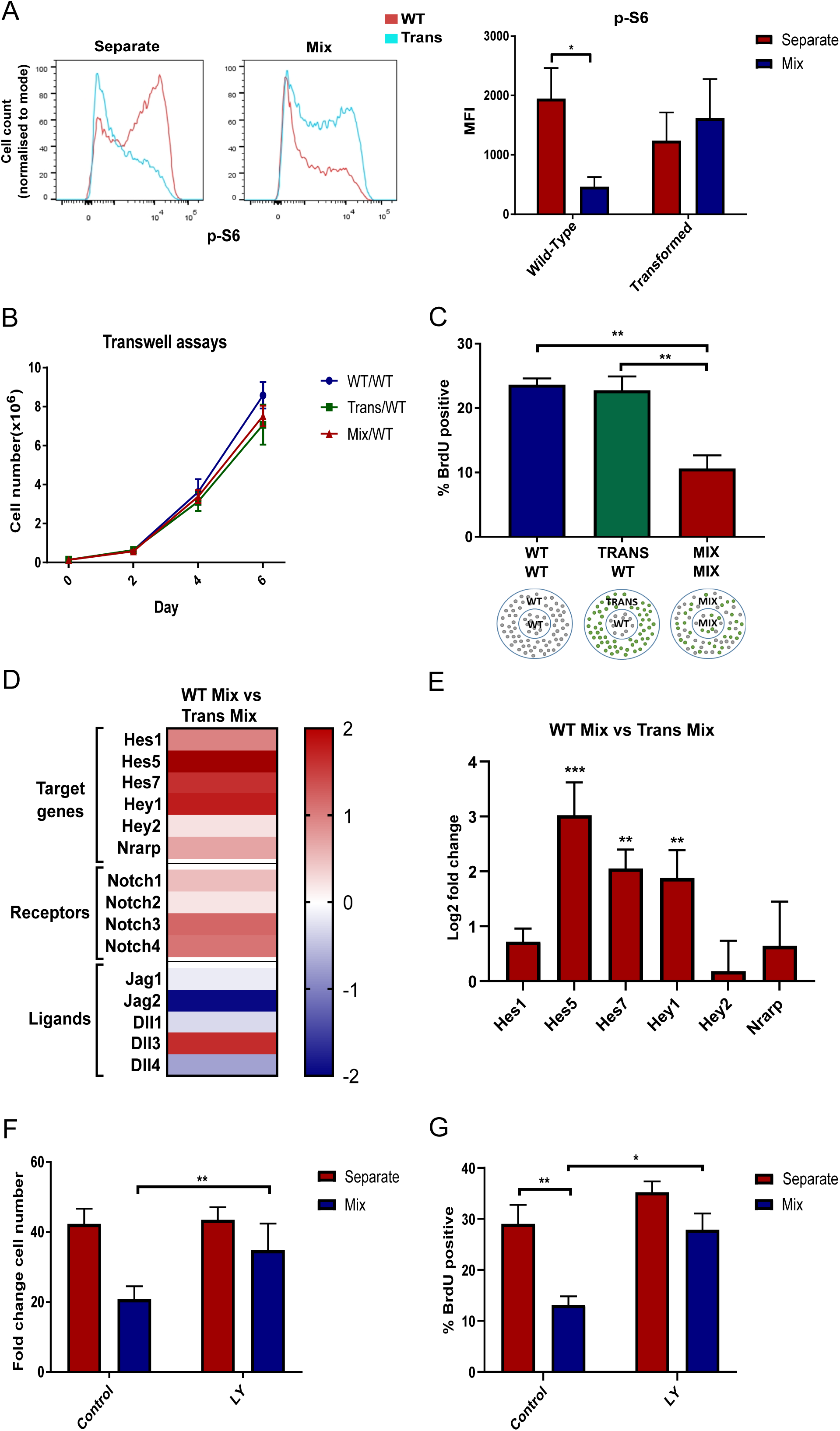
Quiescence of WT NSCs is induced by direct contact with transformed NSCs and is associated with increased Notch activation. **A** Flow cytometry plot showing p-rpS6 staining in WT and transformed NSCs in separate and mixed culture and quantification of median fluorescence intensity. WT NSCs show reduced expression specifically in the co-culture condition. N=6, ANOVA followed by Sidak’s multiple comparisons test. **B** Growth curves of WT NSCs cultured below transwell inserts with WT, transformed or mixed cultures. No difference was observed between conditions. N=3. **C** Percentage of BrdU-positive WT NSCs from the inner ring of the three different cultures shown. When cells were mixed in each well WT NSCs showed a decrease in the proportion of proliferating cells (Mix/Mix). However, WT NSCs surrounded by a ring of transformed NSCs (Trans/WT) did not show a reduction in proliferation relative to WT NSCs surrounded by WT NSCs (WT/WT). N=3, one-way ANOVA followed by Tukey’s multiple comparisons test. Direct cell contact is therefore required for reduced WT NSC proliferation **D** Heat map showing changes in gene expression in co-cultured transformed and WT NSCs. Notch target genes and receptors are upregulated in WT NSCs, while Notch ligands show a more mixed expression with several downregulated relative to transformed NSCs. **E** RT-qPCR validation of Notch target gene expression in WT NSCs relative to transformed NSCs sorted from the co-culture. N=3, two-way ANOVA followed by Sidak’s multiple comparisons test. **F** Fold change cell number quantification of NSC growth with and without addition of LY411575 at 2µM. N=8, two-way ANOVA with Sidak’s multiple comparisons test. **G** Proportion of BrdU positive cells following a 24 hour incubation with LY411575. N=4, two-way ANOVA with Sidak’s multiple comparisons test. WT NSCs in co-culture show increased proliferation in the presence of the γ-secretase inhibitor, LY411575.

Next, we investigated an alternative possibility that cell contact is required to induce the growth arrest of wild-type NSCs. We first used a transwell assay, where the two cell types are cultured in the same well, separated by a permeable membrane that allows the exchange of factors in the media while physically separating the cells. When wild-type cells were co-cultured with transformed cells separated by a transwell, they grew similarly to wild-type cells in homotypic cultures (Figure 3B). This suggests a requirement for cell-cell contact and that secreted paracrine factors are not sufficient to induce the quiescence. We further tested this possibility by using fences, which are metal inserts that divide the well into an inner and outer ring One cell type is plated in the inner ring and the second in the outer ring (Figure 3C) and once they are growing adherently, the inserts can be removed leaving behind a small gap between the two populations. Using this system we compared the proliferation rate of wild-type NSCs that were separated from transformed NSCs to their proliferation when separated from other wild-type NSCs or when mixed with transformed cells. We found that the rate of BrdU incorporation of the wild-type NSCs only decreased when these cells were in physical contact with transformed cells (Figure 3C). This indicates that cell-cell contact is most likely required for transformed cells to inhibit the proliferation of wild-type NSCs.

### Transformed cells signal via Notch and Rbpj to induce quiescence in wild-type NCSs

The Notch pathway is a highly conserved signalling pathway requiring cell-cell contact that plays a central role in the regulation of embryonic and adult neurogenesis (Ables et al., 2011; Urban and Guillemot, 2014). To test the potential role for Notch signalling we first explored the expression of Notch signalling pathway components in our RNA-seq data. Interestingly, genes related to the Notch signalling pathway were enriched in the comparison between co-cultured wild-type and transformed NSCs (p-value = 5.94E^-6^, 21/38 genes differentially expressed with adjusted p-value <0.1). Overall, the majority of Notch-related genes were upregulated in wild-type NSCs compared to transformed NSCs, including Notch targets and receptors, while there was some evidence that Notch ligands were more highly expressed on transformed NSCs in co-culture (Figure 3D). Analysis by real-time qPCR of sorted samples also revealed that the expression of the Notch target genes *Hes1, Hes5, Hes7, Hey1, Hey2* and *Nrarp* are upregulated in co-cultured wild-type NSCs (Figure 3E). This suggests that Notch pathway activity is higher in co-cultured wild-type NSCs than in transformed cells and raises the possibility that transformed cells may be signalling to wild-type NSCs via this pathway.

To test the functional significance of this difference in Notch activation, two complementary γ-secretase inhibitors were used to block Notch signalling. We observed that both LY411575 (LY) and crenigacestat not only reduced Notch target gene expression in NSCs, but also partially rescued the proliferation arrest of wild-type NSCs in co-culture with transformed cells. Evidence for this is that the growth curves and BrdU incorporation rates of co-cultured NSCs treated with the γ-secretase inhibitors were only slightly lower than their growth is separate culture (Figure 3G-H and Supplementary Figure 3A-E). This suggests that Notch signalling is required for the induction of quiescence in wild-type NSCs by transformed cells.

To further test the requirement of Notch signalling in wild-type NSCs we mutated *Rbpj*, a key effector of this pathway, as well as *Notch1* and *Notch2* by CRISPR/Cas9 targeting (Figure 4A, D and G). Importantly, we found that *Rbpj*^*-/-*^, *Notch1*^*-/-*^ or *Notch2*^*-/-*^ NSCs were not susceptible to proliferation arrest when co-cultured with IE-NSCs. For example, we observed that all three Notch pathway mutant cell lines displayed similar growth rates in separate and co-culture (Supplementary Figure 4A-D). Similarly, the fold change in cell number and BrdU incorporation rates of co-cultured *Rbpj*^*-/-*^, *Notch1*^*-/-*^ and *Notch2*^*-/-*^ NSCs was similar to separate culture and significantly higher than that of wild-type NSCs co-cultured with IE-NSCs (Figure 4B-C, E-F and H-I). Interestingly, both the growth curves and final cell counts of *Rbpj*^*-/-*^ NSCs indicated that these cell types even out-grew IE-NSCs in co-culture (Figure 4B and Supplementary Figure 4B), suggesting that disrupting Notch signalling may be sufficient to provide NSCs with a competitive advantage against transformed NSCs. Together, these data indicate that Notch signalling is required for the suppression of proliferation of wild-type NSCs by transformed cells.

**Figure 4.**
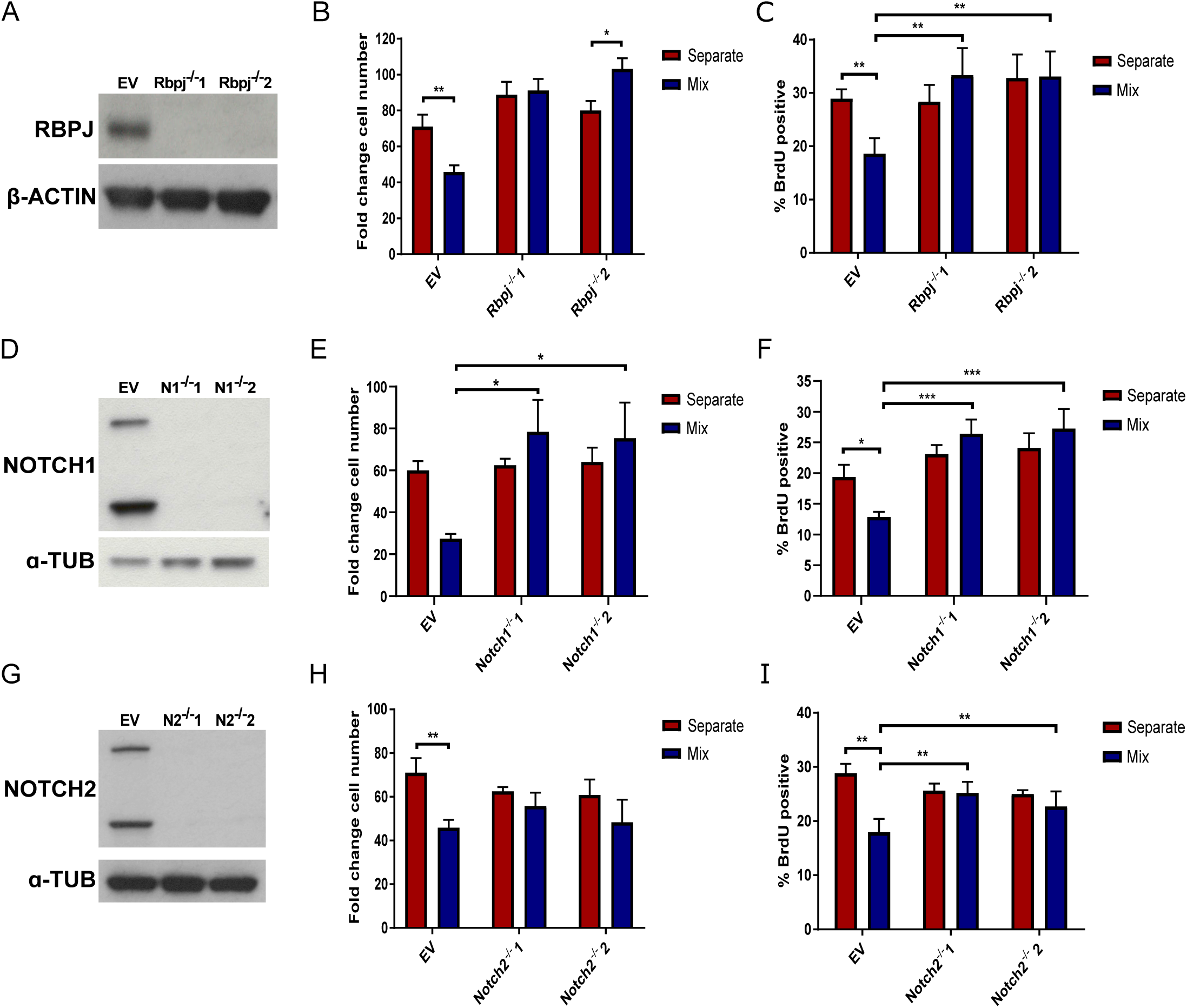
Disruption of Notch signalling rescues WT NSC proliferation in co-culture. **A** Western blot showing absence of RBPJ protein in both *Rbpj*^*-/-*^ clones. **B** Fold change cell number quantification of EV and *Rbpj*^*-/-*^ NSCs in separate and co-culture with transformed NSCs. EV control NSCs show decreased growth in co-culture but this is not the case for the *Rbpj*^*-/-*^ NSCs. N=4, two-way ANOVA followed by Sidak’s multiple comparisons test. **C** Proportion of BrdU-positive cells in separate and co-culture showing decrease in EV control but not in *Rbpj*^*-/-*^ NSCs. N=5, two-way ANOVA followed by Sidak’s multiple comparisons test. **D** Western blot showing absence of NOTCH1 protein in both *Notch1*^*-/-*^ clones. **E** Fold change cell number quantification of EV and *Rbpj*^*-/-*^ NSCs in separate and co-culture with transformed NSCs. EV control NSCs show decreased growth in co-culture but this is not the case for the *Notch1*^*-/-*^ NSCs. N=4, two-way ANOVA followed by Sidak’s multiple comparisons test. **F** Proportion of BrdU-positive cells in separate and co-culture showing decrease in EV control but not in *Notch1*^*-/-*^ NSCs. N=5, two-way ANOVA followed by Sidak’s multiple comparisons test. **G** Western blot showing absence of NOTCH2 protein in both *Notch2*^*-/-*^ clones. **H** Fold change cell number quantification of EV and *Notch2*^*-/-*^ NSCs in separate and co-culture with transformed NSCs. EV control NSCs show decreased growth in co-culture but this is not the case for the *Notch2*^*-/-*^ NSCs. N=4, two-way ANOVA followed by Sidak’s multiple comparisons test. **I** Proportion of BrdU-positive cells in separate and co-culture showing decrease in EV control but not in *Notch2*^*-/-*^ NSCs. N=5, two-way ANOVA followed by Sidak’s multiple comparisons test.

Finally, to exclude the possibility that Notch signalling is required in transformed NSCs, we mutated *Rbpj* in this cell type (Supplementary Figure 4E). We observed that IE-NSCs still induced the proliferation arrest of wild-type NSCs, as shown by the growth curves, final cell numbers and rate of BrdU incorporation (Supplementary Figure 4F-H). Therefore Notch signalling is not required in transformed cells for their growth inhibition of wild-type NSCs.

## Discussion

In conclusion, our data indicate that transformed cells activate Notch signalling in adjacent wild-type NSCs to induce in these a quiescent state. Importantly, the role of Notch signalling in regulating the activated versus quiescent state of NSCs is borne out by the work of others. Increased expression of Notch signalling pathway components in quiescent NSCs compared to activated NSCs has been widely reported in RNA-sequencing studies (Andersen et al., 2014; Llorens-Bobadilla et al., 2015; Morizur et al., 2018). Additionally, NOTCH receptor and RBPJ expression have been previously confirmed in both quiescent and activated NSC populations in the SVZ, and NOTCH2 has been shown to mediate signals promoting NSC quiescence within the niche (Engler et al., 2018). The Notch ligand Dll1 has also been reported to be expressed exclusively in activated NSCs and transit-amplifying cells (Kawaguchi et al., 2013) and its deletion resulted in excessive production of activated NSCs and a depletion of the quiescent NSC pool. Similar observations have also been made in zebrafish, where activated radial glial cells were shown to revert to a quiescent state by Notch pathway activation (Chapouton et al., 2010). Therefore our results imply that transformed cells highjack an evolutionarily conserved pathway regulating vertebrate NSC quiescence and use it to gain a competitive advantage over other NSCs in the niche.

Our work also has implications for glioblastoma tumour progression. The SVZ has been shown to be a permissive region for glioblastoma growth, as tumours in contact with this region are associated with decreased patient survival and increased recurrence (Khalifa et al., 2017; Mistry et al., 2017). This niche has also been suggested to serve as a reservoir for malignant cells away from the tumour mass and may therefore support tumour recurrence (Piccirillo et al., 2015). In humans it has been shown that glioblastoma can arise from SVZ cells with low-level driver mutations (Lee et al., 2018), further highlighting the importance of the SVZ for oncogenic cell propagation. We therefore speculate that the mechanisms we describe operate during early premalignant disease to support tumorigenesis. Here, transformed NSCs provide negative feedback signals to surrounding normal NSCs limiting their self-renewal and enforcing their quiescence. By this mechanism, oncogenic NSCs can ensure their preferential self-renewal and differentiation, therefore increasing both their cell number and their likelihood of giving rise to progenitor cells and exiting the niche. This competitive advantage may be an important early step in the development of glioma and glioblastoma and therefore understanding it may help develop therapeutic approaches.

## Materials and Methods

### Neural Stem Cell Cultures

Adult murine NSCs were derived from the SVZ by the protocol described in [5]. Transformed *Ink4a/Arf*^*-/-*^; EGFRvIII NSCs (IE-NSCs) were generated by the Maarten van Lohuizen laboratory (Netherlands Cancer Institute). NSCs were isolated from *Ink4a/Arf*^*-/-*^ mice and transduced with EGFRvIII pMSCV retrovirus (Bruggeman et al., 2007). All cells were cultured at 37°C in an atmosphere with 5% CO2. NSCs were maintained in serum-free adherent growth conditions on poly-L-lysine-coated plates (Sigma). 1µg/ml Laminin (Sigma) was also added to the growth media to promote adherent growth. Growth medium was Dulbecco’s Modified Eagle Medium: Nutrient Mixture F12 (DMEM/F12) (Gibco) supplemented with N2 (1:200, Invitrogen), B27 (1:100, Invitrogen), 1x non-essential amino acids (Invitrogen), 0.1% BSA (Gibco), 100µM β-mercaptoethanol (Invitrogen), 1.5mg/ml glucose (Sigma), 15mM HEPES (Gibco), 2mM L-glutamine (Gibco), 100U/ml penicillin-streptomycin (Gibco), 20ng/ml EGF and 10ng/ml bFGF (both Peprotech).

### Co-culture assays

For six-day co-culture assays, 40,000 cells per well were seeded in a 24-well plate either separately or as a 50:50 mix of WT and transformed NSCs. Standard growth media was used throughout and was changed on days three, four and five. The proportion of each cell type was assessed by determining the percentage of GFP-positive cells by flow cytometry. Cells were harvested with accutase and resuspended in PBS containing 3% FCS. All samples were transferred into strainer-cap flow cytometer tubes for analysis and 100ng/ml propidium iodide (PI) was added. Flow cytometry was performed on an LSR-II Analyser (BD Biosciences) with FACS Diva software (BD Biosciences). Gates were applied to exclude cellular debris, cell doublets and non-viable PI-positive cells. The percentage of GFP-positive and GFP-negative cells was determined with gates set using separate cell preparations. Separate cultures were also mixed and analysed by flow cytometry. Viable cell counts at each time point were performed using a Vi-Cell XR counter (Beckman Coulter) and calculated for each cell type in separate and co-culture by applying the percentages from the flow cytometry analysis.

For transwell assays, cell culture inserts with 0.4µm pores (Merck Millipore) were placed in a 6-well plate and coated with PLL. 140,000 NSCs were seeded in NSC growth media either onto the insert or below in the well. The growth of WT NSCs seeded below either WT or transformed NSCs or a 50:50 mix was evaluated over six days. For fence assays, 24-well plates were coated with PLL and fences (Aix-Scientifics) were placed in each well. 100,000 NSCs were seeded in the inner ring and 400,000 in the outer ring. The fences were removed 24 hours later and the media was replaced. The following day, 10µM BrdU was added to the wells for two hours. The fences were then replaced into the well and the cells in the inner ring were detached with accutase and harvested. The cells were then fixed in 2% formaldehyde and analysed for BrdU expression by flow cytometry as described below.

### Flow cytometry immunolabelling

Cells were fixed in 2% formaldehyde and permeabilised in ice-cold 90% methanol. Cells were incubated with the primary antibody (anti-p-rpS6, CST 5364, 1:200, anti-BrdU, CST 5292, 1:200) for one hour in PBS with 1% BSA. AlexaFluor 546 secondary antibodies (Molecular Probes) were used for detection) diluted 1:2000 in PBS containing 1% BSA. Secondary incubation was for 30 minutes. Cell debris and doublets were excluded and controls with the primary antibody omitted were used to gate positive and negative cells. Flow cytometry was performed on an LSR-II Analyser (BD Biosciences) with FACS Diva software (BD Biosciences) and data were analysed using FlowJo software.

For BrdU analysis NSCs were incubated for two hours with 10µM BrdU prior to fixation. Before blocking, NSCs were then incubated for 45 minutes at 37°C in DNase I solution consisting of 20µl DNase I (Promega RQ1) in 250µl DNase buffer solution. The immunolabelling protocol was then followed as detailed above.

### FACS

180,000 cells per well were seeded in a 6 well plate either separately or as a 50:50 mix of WT and transformed NSCs and cultured under the standard assay conditions until day five. GFP-negative WT NSCs were sorted using a FACS Aria cell sorter (BD Biosciences) from both separate and mix samples. Gates were applied to exclude cell debris and doublets. Cells were recovered into growth media at room temperature. Cells were then either lysed for RNA/protein extraction or, for replating experiments, counted and replated into a 24-well plate at 100,000 cells per well.

### Western blot analysis

Cells were lysed directly in Laemmli buffer (50mM Tris-HCl (pH 6.8), 2% SDS, 10% glycerol in ddH_2_O) and heated at 95°C for 10 minutes. Protein quantification was carried out using a BCA Protein Assay kit (Pierce) according to the manufacturer’s instructions. 15-20μg denatured protein from each sample was run in XT sample buffer (BioRad) with 0.25% β-mercaptoethanol on 10% or 12% SDS-PAGE Bis-Tris gels (pre-cast BioRad). Proteins were then blotted onto PVDF membranes (Thermo Fisher Scientific) using vertical wet transfer at 100V for one hour at 4°C. Blocking was performed in 5% milk in TBST buffer and primary antibody incubation was done overnight at 4°C in TBST containing 5% BSA. The following antibodies were used at the stated concentration: anti-α-tubulin (CST 3873, 1:1000), anti-β-actin (CST 5970, 1:1000), anti-RBPJ (CST 5313, 1:1000), anti-Notch1 (CST 3608, 1:1000), anti-Notch2 (CST 5732, 1:1000), anti-p-SMAD1/5/8 (CST 9511, 1:000), anti-TSC2 (CST 4308, 1:1000). Horseradish peroxidase-conjugated secondary antibodies (Abcam) diluted 1:5000 in 5% milk in TBST were used for detection by the addition of an enhanced chemiluminescence (ECL) substrate (Promega). Western blot quantification was performed using Fiji software (Schindelin et al., 2012).

### RNA Extraction and Quantitative RT-PCR

RNA was extracted with the RNeasy mini kit (Qiagen) and SuperScript III reverse transcriptase (Thermo Fisher Scientific) was used for cDNA synthesis according to manufacturer’s instructions. RT-qPCR was performed using SYBR-Green Mastermix with ROX reference dye (Sigma). A list of all primer sequences used can be found in Table 1 below. RT-qPCR was performed on an ABI 7900HT Fast Real Time PCR machine (Applied Biosystems) and analysed on SDS Biosystem software. Relative gene expression was calculated using the ΔΔCt method normalised to β-actin expression (Livak and Schmittgen, 2001).

### RNA-sequencing analysis

Sorting was performed by FACS at day five of the six-day assay as described above for three biological repeats. PI was added to exclude non-viable cells. Cells were recovered into growth media and then resuspended in RLT lysis buffer (Qiagen). RNA extraction was performed using the Qiagen RNeasy kit according to the manufacturer’s instructions. RNA samples were quantified using a Qubit fluorometer (Thermo Fisher Scientific) and the quality assessed by TapeStation electrophoresis (Agilent). mRNA was isolated using oligo dT beads. mRNA was then fragmented, converted to cDNA and ligated to Illumina adapters. Following sample indexing, the quality of cDNA libraries was also assessed by TapeStation. Sequencing was performed using the HiSeq 4000 system (Illumina). Sequencing reads were aligned to the mouse genome (mm9) using TopHat2 (Kim et al., 2013) and differential expression was analysed using the DESeq2 package (Love et al., 2014). The resulting gene sets were analysed using Ingenuity Pathway Analysis (IPA) software (Qiagen) (Kramer et al., 2014). This utilises the Ingenuity Knowledge Base for functional annotations and interactions of genes. A ‘Core Analysis’ was performed for each comparison on genes differentially expressed with a padj cut-off <0.1, corresponding to a false discovery rate (FDR) of 10%. This identified enriched Canonical Pathways and Upstream Regulators in each comparison.

### Generation of *Rbpj*^*-/-*^, *Notch1*^*-/-*^, *Notch2*^*-/-*^ and *Tsc2*^*-/-*^ NSCs

gRNAs for targeting of *Rbpj, Notch1* and *Notch2* were cloned into the pX330-U6-Chimeric_BB-CBh-hSpCas9 (PX330) expression plasmid (gift from Feng Zhang, Addgene plasmid #42230) using the protocol provided by the Zhang laboratory (Ran et al., 2013). gRNAs for targeting *Tsc2* were cloned into gRNA expression plasmids (gift from George Church, Addgene plasmid #41824) (Mali et al., 2013). These were then co-transfected with an hCas9 plasmid (gift from George Church, Addgene plasmid #41815). gRNA sequences can be found in Supplementary Table 2. NSCs were transfected with plasmids by nucleofection (AMAXA 2B, Lonza) using programme A-033. NSCs were harvested with Accutase and 2-3×10^6^ cells were resuspended in 100µl mouse neural stem cell nucleofector buffer (Lonza) with 2µg plasmid DNA. NSCs were either transfected with 2ug PX330 vector containing gRNAs targeted to *Rbpj, Notch1* or *Notch2* or 0.6µg each of hCas9 and the two gRNA expression plasmids containing the *Tsc2*-targeting gRNAs. NSCs were also co-transfected with 0.125ug of Puro-pPyCAGIP vector or linear hygromycin marker (Clontech 631625) for selection. Empty vector or Cas9 only controls were also performed. Cells were recovered post-transfection with pre-warmed NSC growth media into a 10cm^2^ dish. Selection with 0.25ug/ml puromycin or 250ug/ml hygromycin was performed from 48 hours post-transfection until the emergence of resistant colonies. These were then picked manually with a p20 pipette and transferred to a 48 well plate for expansion and analysis.

## Quantification and Statistical Analysis

Statistical analysis and data representation were performed using GraphPad Prism software. Statistical methods used are indicated in the relevant figure legends. No randomisation or blinding was used in experiments. Sample sizes were selected based on the observed effects and listed in the figure legends. The sample size (n) is described in the figure legends and refers to the number of independent replicate experiments performed. Adjusted p values are displayed as * p<0.05, ** p<0.01 and *** p<0.001.

## Data and Code availability

The raw data for the RNA-sequencing of wild-type and transformed NSCs are being deposited in the arrayexpress database and an accession code that will be made available prior to acceptance. The authors declare that all data supporting the findings of this study are available within the article and its supplementary information files or from the corresponding author upon reasonable request.

## Key Resources Table

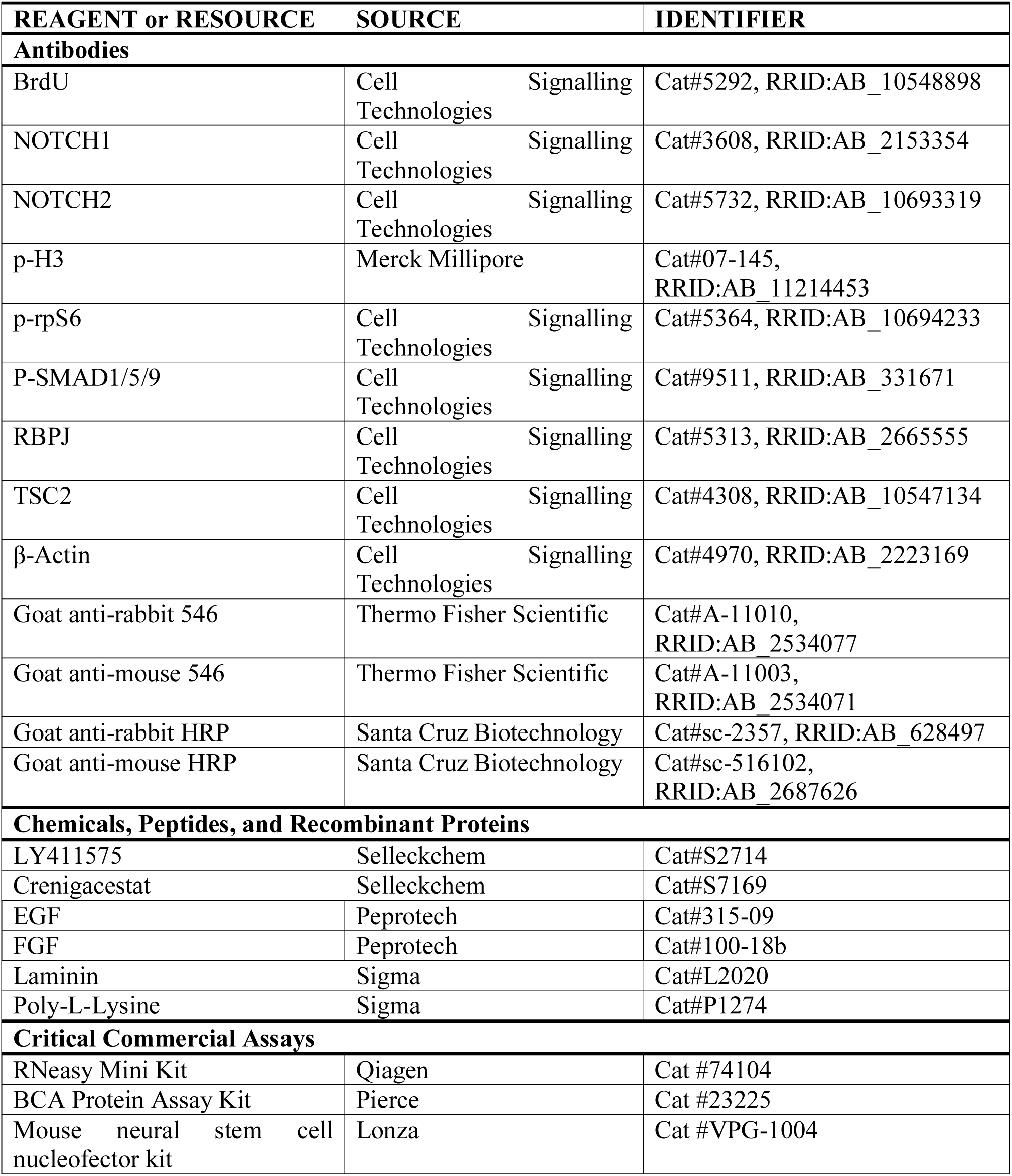

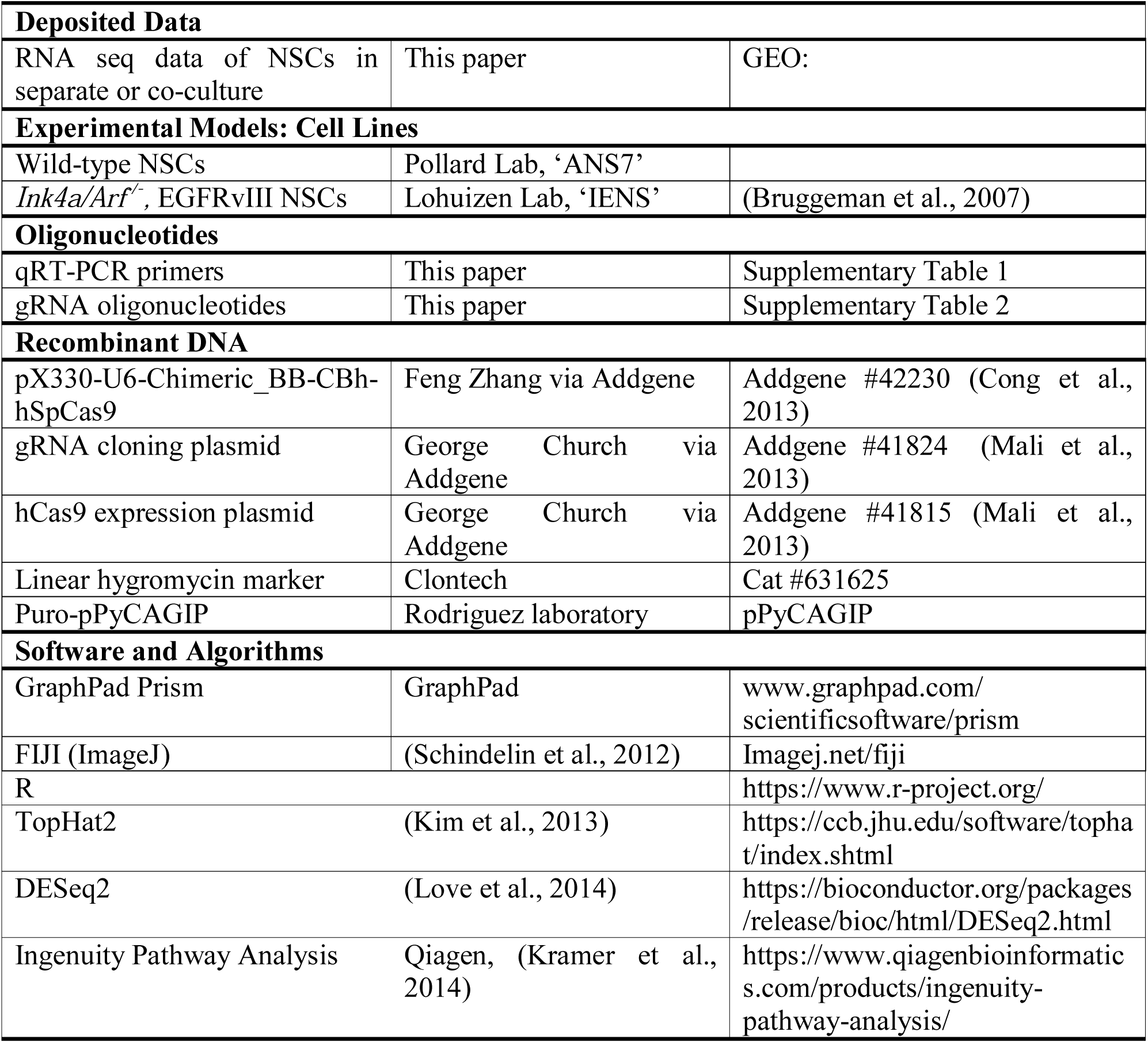

## Acknowledgments

We would like to thank Stephen Rothery for guidance and advice with confocal microscopy. Gratitude also goes to James Elliot for performing cell sorts. Research in Tristan Rodriguez lab was supported by the MRC project grant (MR/N009371/1), by the Rosetrees Trust grant (M693) and by the British Heart Foundation centre for research excellence. Katerina Lawlor was supported by an NHLI PhD studentship.

## Figure Legends

**Supplemental Figure 1.**
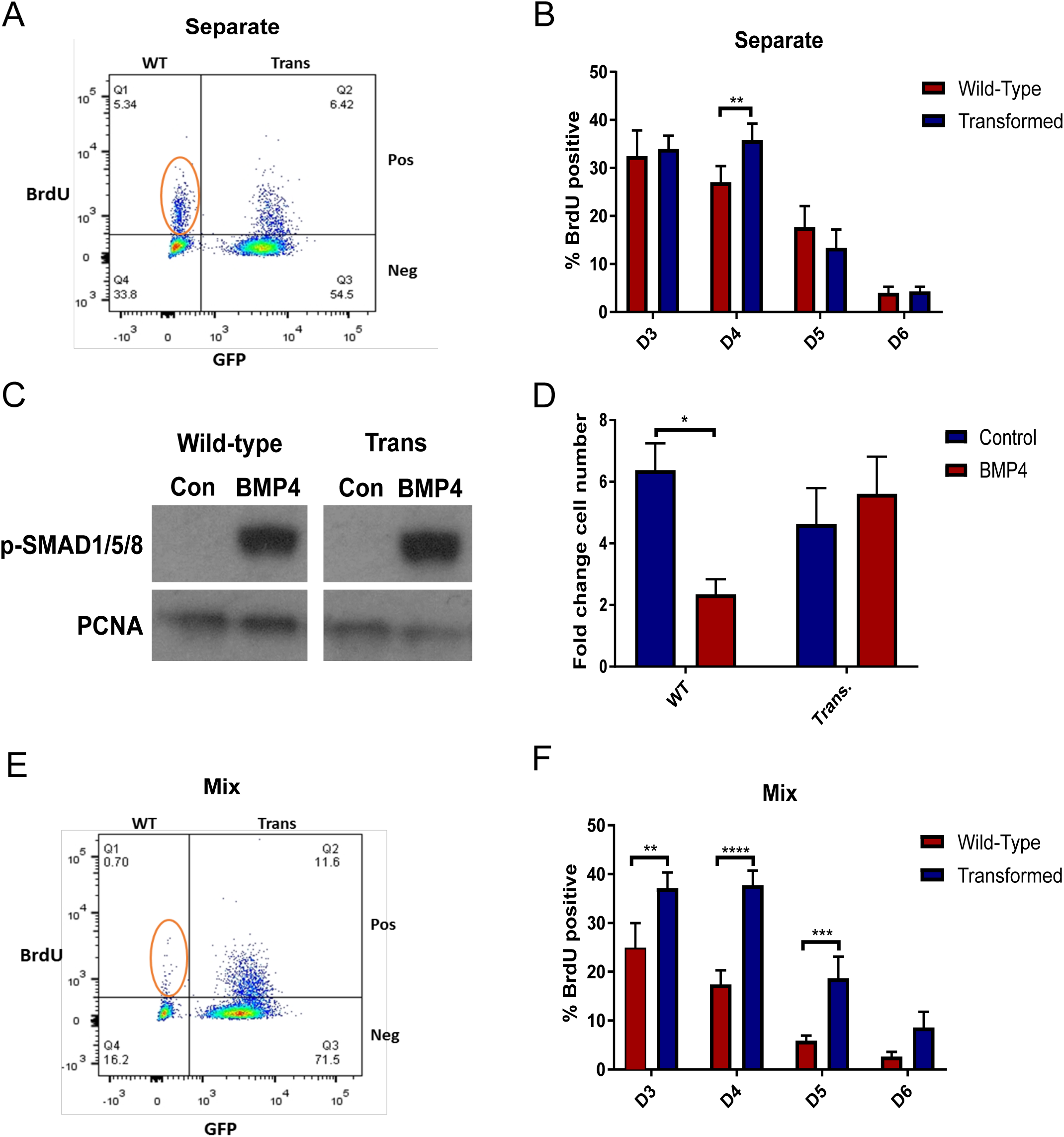
**A** Representative flow cytometry plot of BrdU staining in separately cultured NSCs at day five. **B** Time-course of BrdU staining throughout the assay in separate culture. N=5, ANOVA followed by SIdak’s multiple comparisons test. **C** Fold change cell number quantification of NSCs cultured in the presence or absence of 50ng/ml BMP4. BMP4 treatment reduces the growth of WT, but not transformed, NSCs. N=4, two-way ANOVA followed by Sidak’s multiple comparisons test. **D** Western blot showing increased p-SMAD1/5/8 expression in NSCs treated with 50ng/ml BMP4 for 24 hours relative to control NSCs (Con). **E** Representative flow cytometry plot of BrdU staining in co-cultured NSCs at day five. **F** Time-course of BrdU staining throughout the assay in co-culture showing reduction in BrdU incorporation in WT NSCs. N=5, ANOVA followed by Sidak’s multiple comparisons test.

**Supplemental Figure 2.**
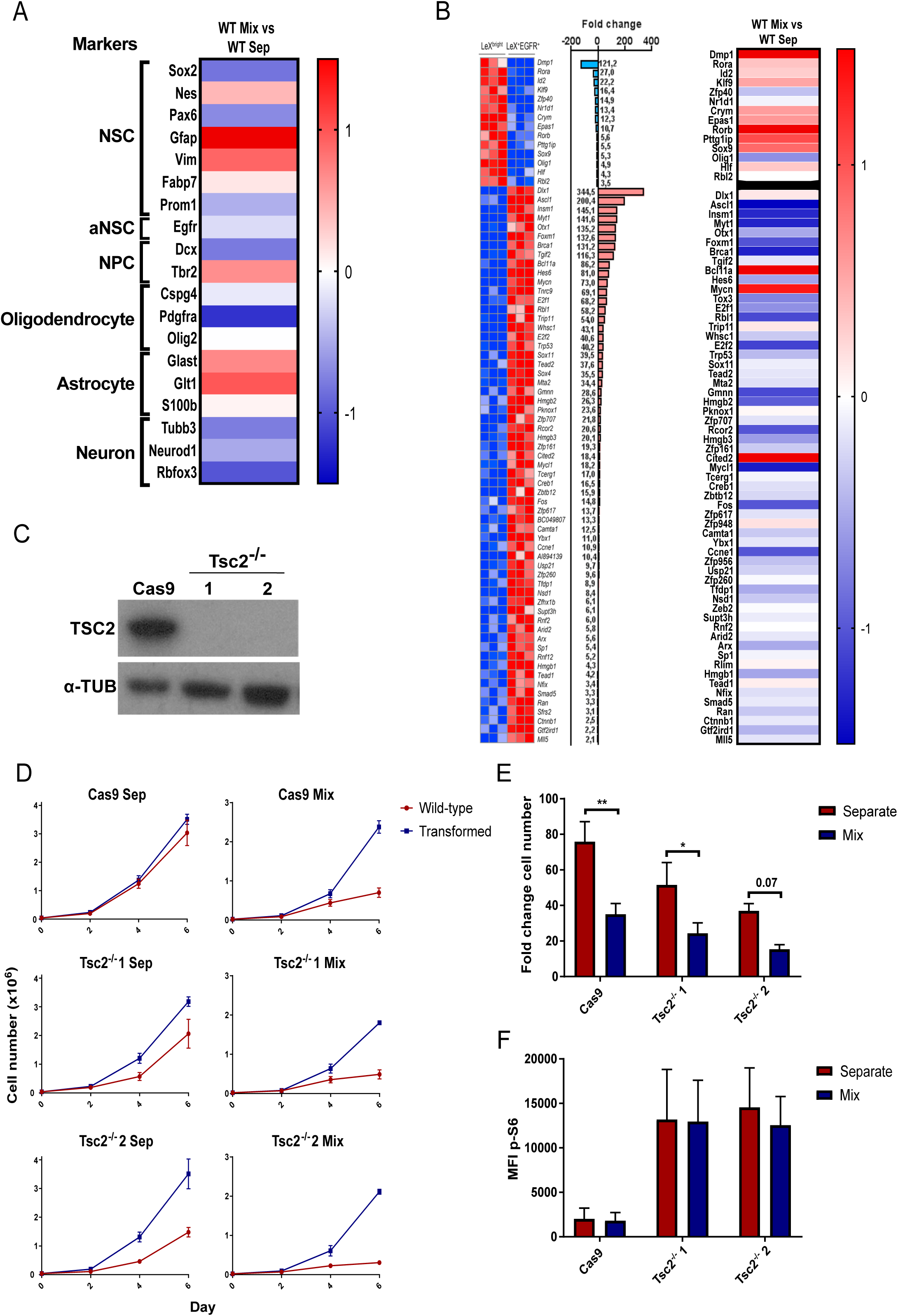
**A** Heat map showing changes in gene expression in the ‘WT Mix vs WT Sep’ comparison. Classical marker genes for different cell populations are shown including those used commonly to identify adult NSCs. GFAP log_2_ fold change is off the scale at 5.4. **B** Comparison to published transcriptional analysis of quiescent and activated NSCs. Left - Transcription factors and co-factors differentially expressed in quiescent and activated NSCs from [18]. Right – Heat map showing the expression of the genes identified by Morizur et al in the comparison of WT NSCs in separate (WT Sep) or co-culture (WT Mix). Genes enriched in quiescent NSCs were broadly upregulated in co-cultured WT NSCs, while those enriched in activated NSCs were broadly downregulated. **C** Western blot showing absence of TSC2 expression in *Tsc2*^*-/-*^ clones. The control Cas9 clone was transfected with Cas9 only without the gRNAs. **D** Growth curves of transformed NSCs and Tsc2^-/-^ or Cas9 control NSCs in separate or co-culture. *Tsc2*^*-/-*^ clones grow more slowly than control and still show reduced growth in co-culture. N=4. **E** Fold change cell number quantification of growth curves. N=4, two-way ANOVA followed by Sidak’s multiple comparisons test. **F** Expression of p-rpS6 in Cas9 and *Tsc2*^*-/-*^ clones as determined by flow cytometry. mTOR activation is increased in *Tsc2*^*-/-*^ NSCs and remains high in co-culture despite reduced growth. N=3.

**Supplemental Figure 3.**
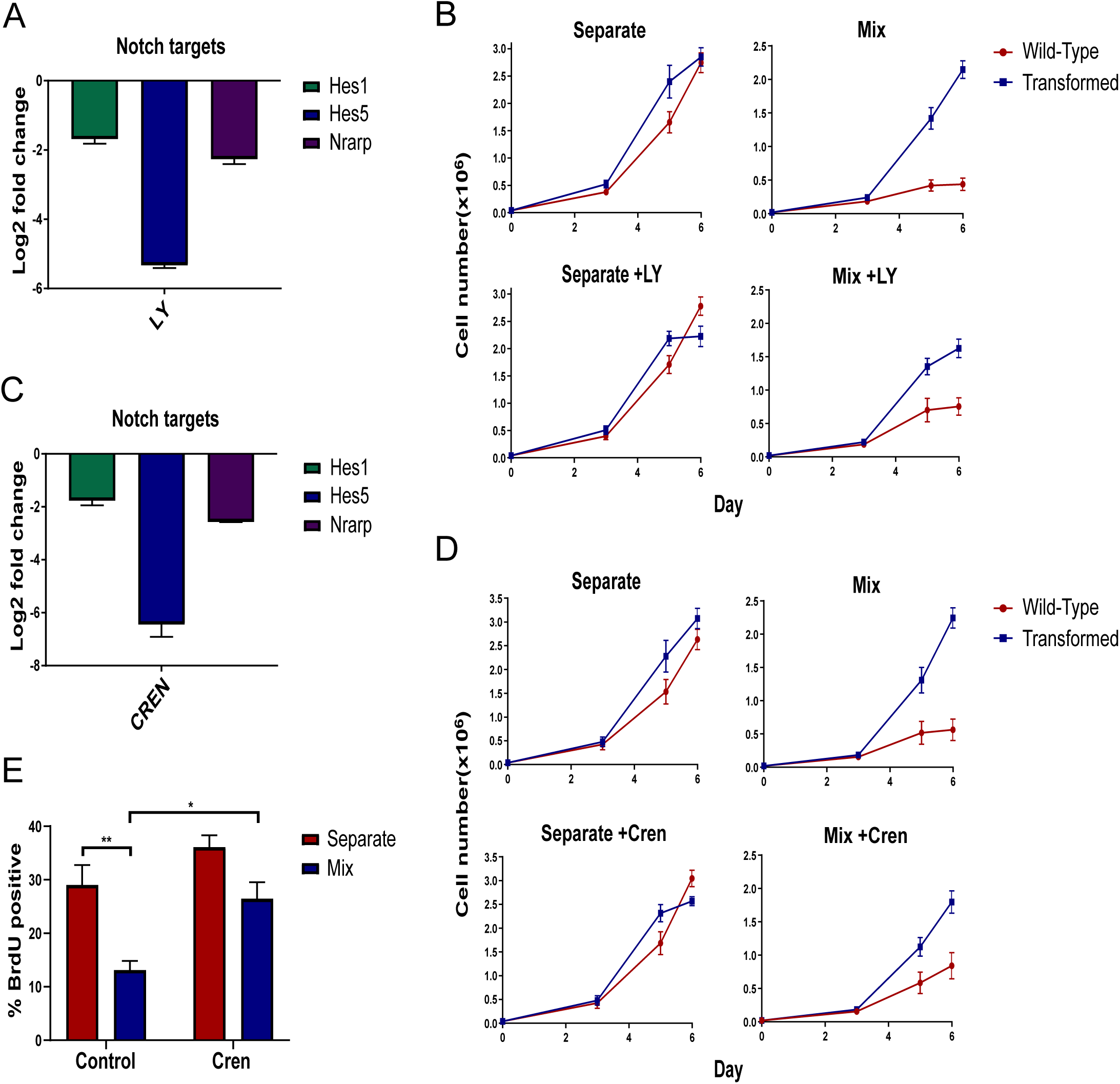
**A** RT-qPCR analysis of the Notch target genes, *Hes1, Hes5* and *Nrarp* in WT NSCs treated with 2µM LY411575. Log_2_ fold change relative to control NSCs. N=3. **B** Growth curves of WT and transformed NSCs cultured in the standard assay with or without LY411575 addition from day three. **C** RT-qPCR analysis of the Notch target genes, *Hes1, Hes5* and *Nrarp* in WT NSCs treated with 1µM crenigacestat. Log_2_ fold change relative to control NSCs. N=3. **D** Growth curves of WT and transformed NSCs cultured in the standard assay with or without crenigacestat addition from day three. N=8, two-way ANOVA followed by Sidak’s multiple comparisons test. **E** Proportion of BrdU positive cells following a 24 hour incubation with crenigacestat. N=4, two-way ANOVA with Sidak’s multiple comparisons test. The γ-secretase inhibitor crenigacestat also increases WT NSC proliferation in co-culture.

**Supplemental Figure 4.**
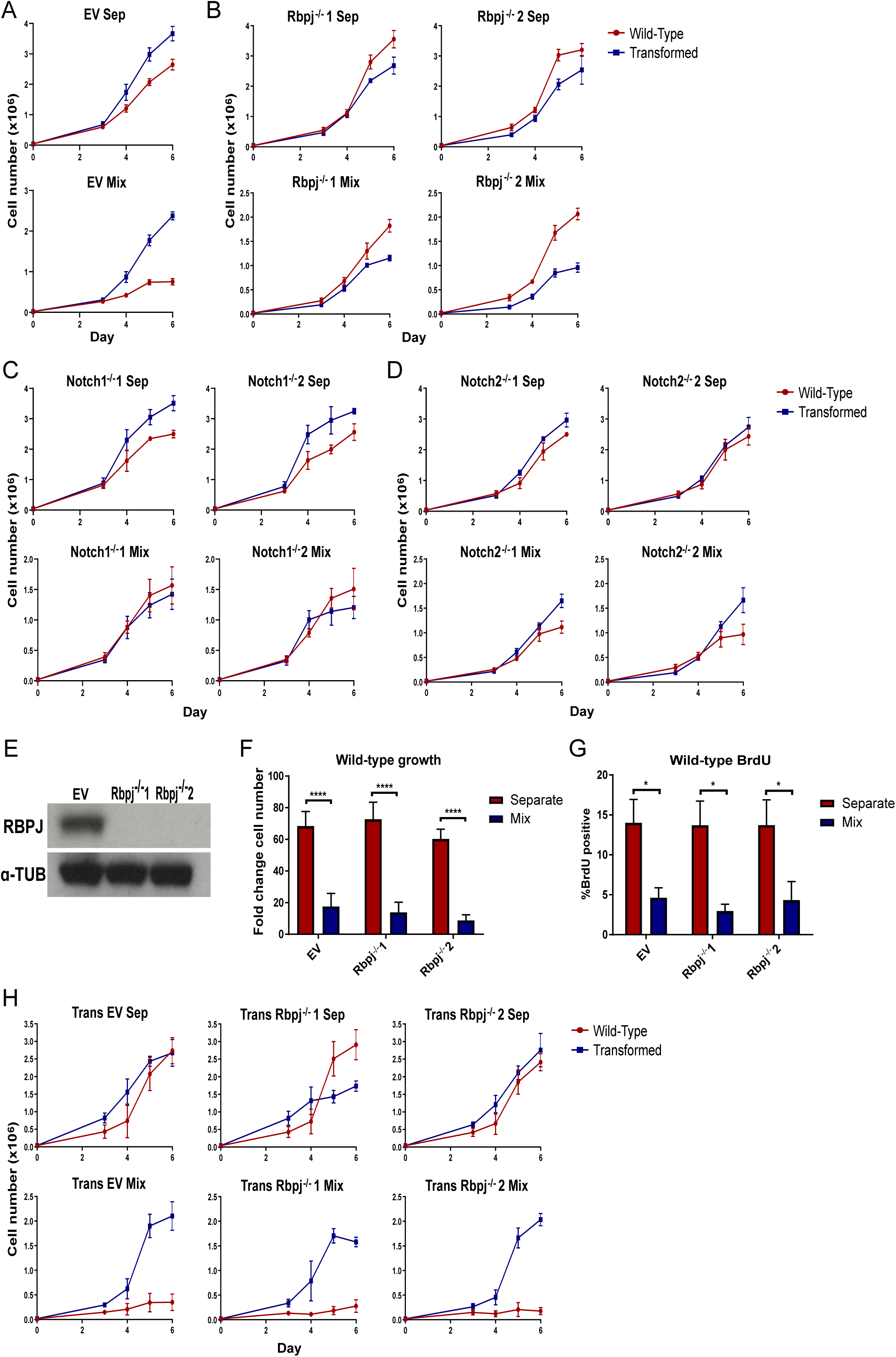
**A** Growth curves of control EV clone cultured separately or in co-culture with transformed NSCs. **B** Growth curves of *Rbpj*^*-/-*^ clones cultured separately or in co-culture with transformed NSCs. **C** Growth curves of *Notch1*^*-/-*^ clones cultured separately or in co-culture with transformed NSCs. **D** Growth curves of *Notch2*^*-/-*^ clones cultured separately or in co-culture with transformed NSCs. N>4. **E** Western blot showing absence of RBPJ protein from *Rbpj*^*-/-*^ transformed NSCs. **F** Fold change cell number of WT NSCs in separate or co-culture with EV or *Rbpj*^*-/-*^ transformed NSCs. N=5, two-way ANOVA followed by Sidak’s multiple comparisons test. **G** Proportion of BrdU-positive WT NSCs in separate or co-culture with EV or *Rbpj*^*-/-*^ transformed cells. N=3, two-way ANOVA followed by Sidak’s multiple comparisons test. **H** Growth curves of transformed EV and *Rbpj*^*-/-*^ clones cultured separately or in co-culture with WT NSCs. Both *Rbpj*^*-/-*^ clones suppress the proliferation of WT NSCs to a comparable degree to the EV control transformed cells.

**Supplementary Table 1.**
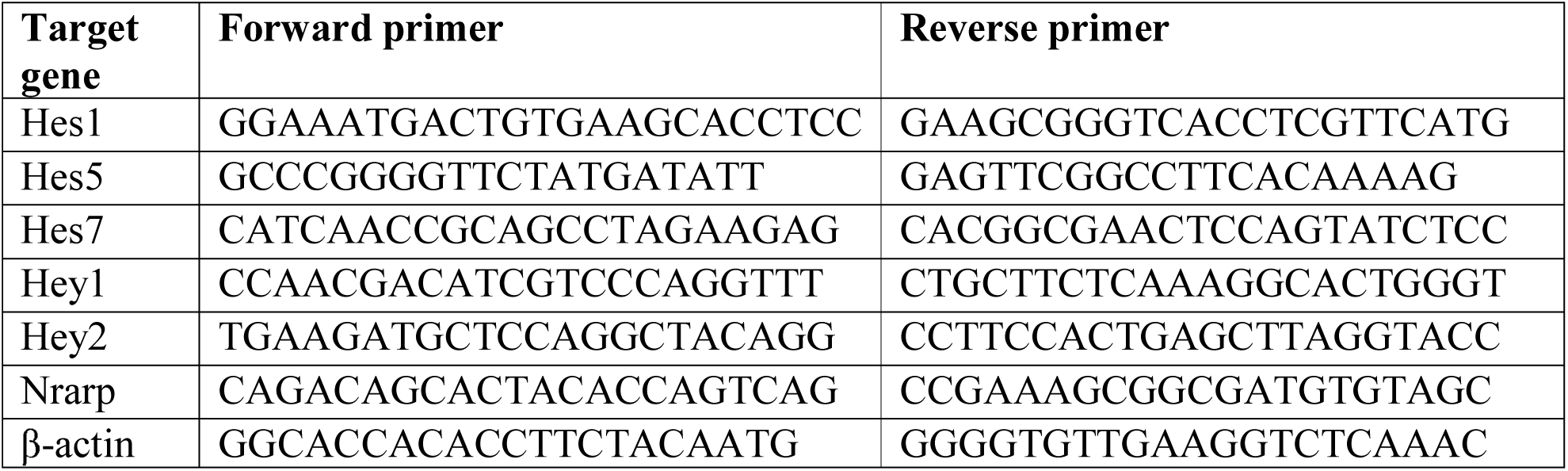
Primers used in qPCR.

**Supplementary Table 2.**
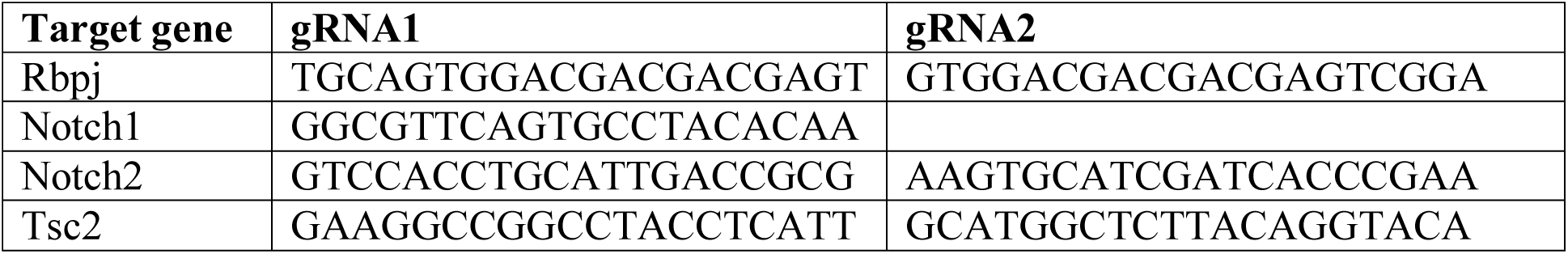
gRNA sequences for deletion.

